# Silent Speech Recognition with Wearable Magnetometers

**DOI:** 10.1101/2025.08.04.668236

**Authors:** Debadatta Dash, Evan Kittle, Isabel Gerrard, Richard Csaky, Gabriel Gonzalez, David Taylor, Juan Pablo Llinas, Dominic Labanowski, Nishita Deka, Richy Yun

**Affiliations:** Sonera Inc.

## Abstract

Next-generation human-computer interaction (HCI) is moving towards more seamless, intuitive, and personal modes of communication, redefining how we interact with technology and one another. Within this landscape, silent speech recognition (SSR) offers a powerful new interaction paradigm, enabling hands-free, private interaction while supporting individuals with speech impairments and enabling communication in noisy or sensitive environments. Recent advances in miniaturized sensors and artificial intelligence (AI) have accelerated the development of more sophisticated wearable SSR systems, driven by growing demand for effortless and accessible communication. Although electrophysiological (ExG) modalities, particularly electromyography (EMG), have dominated early efforts in developing wearable SSR, critical challenges remain. Limited generalizability across users, sensor-skin interface issues, and difficulties with the comfort of use are all current roadblocks to reliable, high-fidelity signals in a wearable form factor. We propose that magnetometers offer a promising alternative to ExG and have the potential to unlock more robust, generalizable, and user-friendly SSR systems. We demonstrate that magnetometers embedded in a headphone form factor achieve a per-user SSR accuracy of 86%, significantly outperforming previously reported state-of-the-art wearable headphones combining ExG and inertial measurement units (IMUs). In addition, we show that wearable magnetometry enables generalization across individuals for SSR. Extending beyond headphones, we also introduce a necklace form factor with magnetometers that is capable of decoding both silent and overt speech in ambient conditions, further showcasing the versatility of magnetometers across different wearable designs in real-world conditions.

## 1 Introduction

The rapid evolution of personal computing and human-computer interaction (HCI) continues to drive the demand for more intuitive, seamless, and context-aware communication modalities. As wearable technologies become increasingly embedded in daily life, there is a growing need to explore new input methods that move beyond traditional touch-based interfaces. Among these, speech as an input modality is gaining significant traction, due to its ease of use, speed, and hands-free approach. Whether vocalized or silent, speech offers key advantages over manual input methods like typing or touch, particularly in mobile, multitasking, or accessibility-focused contexts (Ruan et al., 2018). It supports a wide range of users and can function effectively even when visual or tactile attention is limited.

Within this evolving landscape, silent speech recognition (SSR) has emerged as a compelling alternative to traditional voice input. SSR—the ability to decode speech by interpreting subtle articulatory cues without relying on audible vocalization (Denby et al., 2010; Meltzner et al., 2017)—provides critical communication support for individuals with speech impairments (Cao et al., 2021; Gonzalez-Lopez et al., 2020; Meltzner et al., 2017), offers enhanced privacy in noisy or sensitive environments (Pandey et al., 2021), and unlocks new modes of hands-free HCI (Freitas et al., 2014; Kimura et al., 2022; Sahni et al., 2014). These capabilities are especially relevant in today’s increasingly mobile and connected world, where seamless, context-aware communication is becoming ever more essential (Kimura et al., 2022; Pandey et al., 2021). The rapid advancements and widespread adoption of wearable technologies have further catalyzed interest in integrating SSR into wearable silent speech interfaces (SSIs) (Cao et al., 2023; Gao et al., 2020; Igarashi et al., 2022; Jin et al., 2022; Kapur et al., 2018; Srivastava et al., 2024; Tang et al., 2024). Over the past decade, a diverse set of SSIs has been developed to realize this vision, leveraging a wide range of sensing modalities to capture and interpret non-vocalized speech by tracking articulator movements such as the lips, tongue, and jaw (Freitas et al., 2017). Early efforts largely centered on visual methods, leveraging cameras and computer vision techniques to detect lip shapes and facial motions associated with silent articulation (Hueber et al., 2008; K. Sun et al., 2018). While effective in controlled settings, visual methods often require a clear line of sight and are sensitive to lighting conditions, limiting their practical use in wearable systems. Alternative approaches have included wireless signal perturbations (Birkholz et al., 2018; Khanna et al., 2019) and RFID-based techniques (Wang et al., 2019), where disruptions in signal patterns caused by articulator movements were used to infer speech content. Other approaches in the sensing landscape for SSIs have further diversified: ultrasound imaging has enabled detailed tracking of tongue motion (Hueber et al., 2008; Kimura et al., 2019), but requires bulky hardware; acoustic sensing (Fu et al., 2024; Luo et al., 2021) has captured subtle sounds produced even during silent articulation but remains vulnerable to external noise; and inertial measurement units (IMUs) (Kwon et al., 2023) have been used to detect gross movements of the jaw and lips, but cannot capture fine-grained motion.

In contrast to these approaches, physiological sensing captures biosignals at the source of articulation, directly measuring the neuromuscular activity that drives silent speech, offering a more robust approach to SSR. Unlike vision- or acoustic-based methods, physiological sensing is inherently less sensitive to environmental factors, making it especially promising for wearable and mobile applications. Among physiological sensing modalities, electrophysiological sensing (ExG; electromyography (EMG), electroencephalography (EEG), electrooculography (EOG), etc.), particularly surface electromyography (sEMG), has led much of the early work in wearable SSR (Meltzner et al., 2018; Song et al., 2023; Zhu et al., 2020). A recent study (Srivastava et al., 2024) proposed a novel multi-modal system (QuietSync) that combined IMUs and ExG electrodes to enable SSI across four different form factors—headphones, earphones, glasses, and VR/XR systems. Among the proposed form factors, the headphone form factor (with ExG and IMUs around the ears) was the most robust, achieving about 80% average classification accuracy across 9 users over 12 commands (e.g., “Call Mom”) in a user-dependent setting (i.e., a SSR model is trained and evaluated on data from the same user). Importantly, performance improved by more than 10% when IMUs were combined with ExG signals compared to ExG alone, highlighting the challenges of relying solely on ExG for wearable SSR. Similarly, another study focused on overt speech (Wu et al., 2024), classified 11 similar articulatory words (e.g., “aga”, “aba”) with 92.7% user-dependent accuracy using sEMG channels in a necklace form factor. Decoding overt speech benefits from additional signal sources such as glottal activity, airflow, and acoustic cues (Srivastava et al., 2024), which are significantly attenuated or absent during silent speech production. This fundamental difference highlights the unique challenge of silent speech recognition and further motivates the search for sensing modalities that can robustly capture subtle articulatory cues under these constraints.

Despite promising results, ExG-based wearable systems, including sEMG, face several additional limitations that hinder their effectiveness for wearable silent speech recognition: the requirement of skin-contact, leading to electrical signal distortion due to tissue impedance, and limited generalizability due to variability in electrode placement, skin condition, and user-specific anatomical differences (Deng et al., 2014; Yun, Gonzalez, et al., 2024; Zhang et al., 2025). As a result, there is a pressing need for next-generation physiological sensing modalities that can retain the benefits of direct neuromuscular measurement while enhancing robustness, wearability, and generalizability for practical SSIs.

Magnetometers offer an exciting new frontier for wearable SSIs. Instead of solely relying on the electrical signals from muscles at the skin’s surface, magnetometers can measure both mechanical muscle vibrations (mechanomyography) and electrical neuromotor activity (magnetomyography, MMG) (Cohen & Givler, 1972; Yun, Gonzalez, et al., 2024), together defined here as MxG. This approach does not need skin contact, making it inherently less sensitive to surface variability and changes in tissue impedance (Elzenheimer et al., 2020; Yun, Csaky, et al., 2024). Magnetometers have already proven their value in various human-computer interaction settings — from gesture recognition (Jonna & Rao, 2023; Lee et al., 2016; Meier et al., 2019; Yun, Csaky, et al., 2024) to motion tracking (Ismail et al., 2019; Woodward et al., 2014). In this study, we investigated SSR with form factors (headphone and necklace) similar to previous studies with ExG (Srivastava et al., 2024; Wu et al., 2024), but with magnetometers instead of electrical sensors. We found that wearable magnetic sensing enables SSR performance that rivals and potentially exceeds existing state-of-the-art approaches.

## 2 Results

To evaluate the potential of magnetometers for wearable SSR, we collected MxG data from 9 participants (users) using 20 optically pumped magnetometers (OPMs) embedded into a pair of custom 3D-printed over-ear headphones (Figure 1B). Data collection was conducted in a magnetically shielded room (MSR) while participants silently articulated 12 distinct commands or phrases. Each trial was of 1.5 seconds duration with a 1.5 second break between trials and was repeated 40 times per command per session (Figure 1A).

**Figure 1:**
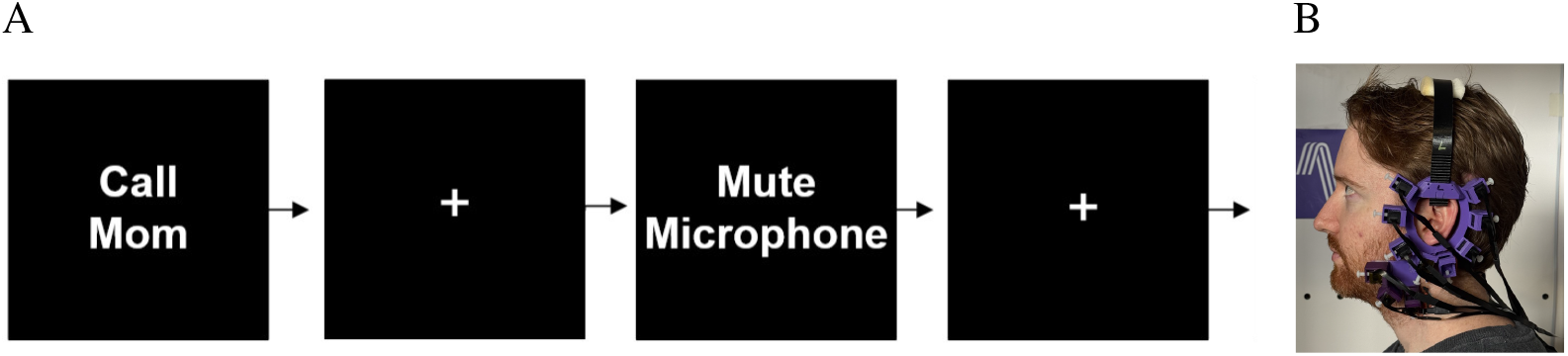
Experimental Setup. **(A) Silent speech task**. Participants silently articulated 12 commands as in (Srivastava et al., 2024). These were: ‘Call Mom’, ‘Clear Notification’, ‘Clear Calendar’, ‘Lock Screen’, ‘Close Camera’, ‘Open Camera’, ‘Mute Microphone’, ‘Join Meeting’, ‘Leave Call’, ‘Decrease Volume’, ‘Increase Volume’, and ‘Play Music’. Each command was displayed on the screen for 1.5 s, followed by a rest (neutral) period of 1.5 s, where a fixation cross replaced the command. When a command was displayed on the screen, the participants performed the silent speech task, i.e., silently articulated the command without any audible voice. Each command was repeated for 40 trials in a pseudo-randomized order. **(B) Headphone form factor for data collection**. Headphone worn by one of the authors for demonstration. Ten two-axis (radial and tangential) QZFM Gen-2 optically pumped magnetometers (OPMs) were embedded in custom 3D printed headphones on each side of the ear for a total of 20 sensors. The experiment took place in a magnetically shielded room (MSR), with the projector placed outside the MSR displaying the stimuli.

### 2.1 Magnetometers in a headphone form factor surpass state-of-the-art SSR performance

We assessed user-dependent (training and evaluating a model on data from the same user) SSR performance by classifying the MxG signals corresponding to the 12 silent commands (Figure 2A). Using an 8-fold cross-validation strategy within each participant, we evaluated performance using 26 different linear classifiers. Ridge classifier consistently achieved strong cross-validation performance (Supplementary Figure S1), resulting in an average classification accuracy of 85.95% (SD = 6.56%) across the 9 participants (Figure 2). Individual accuracy ranged from 77.1% to 95.7%, with 8 out of 9 participants achieving over 80% accuracy. Furthermore, 9 of the 12 commands exhibited over *∼* 80% sensitivity (Figure 2B). Most misclassifications occurred between phonetically similar commands such as “**i**ncrease volume” and “**d**ecrease volume,” which differ by only a single phoneme. The performance achieved with magnetometers significantly surpassed the previously reported ~ 80% classification accuracy for the same 12 commands using combined ExG and IMU sensors in a headphone form factor (Srivastava et al., 2024), highlighting magnetometers as a potentially superior solution for high-fidelity wearable silent speech recognition.

**Figure 2:**
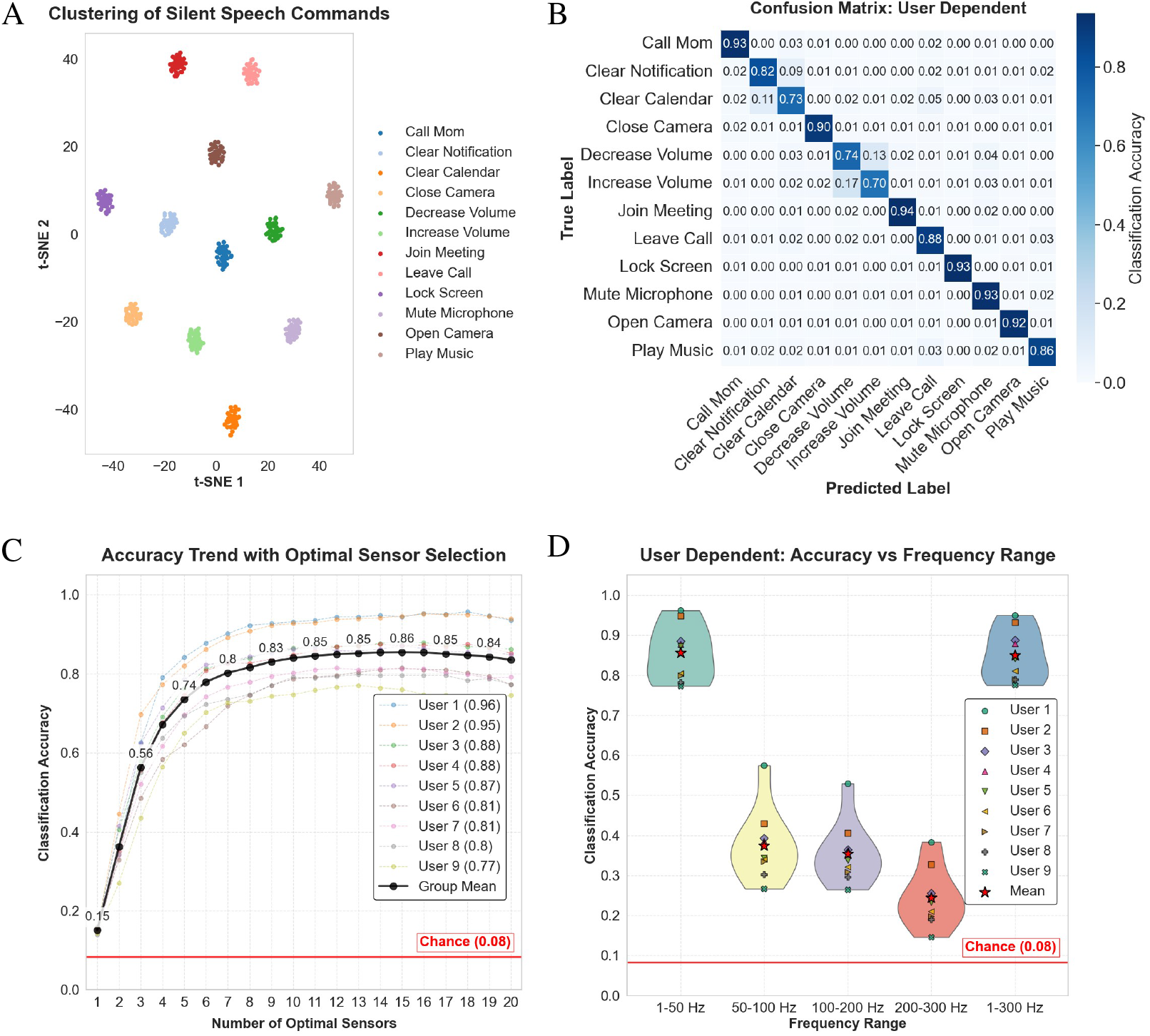
User-dependent SSR performance with headphone-MxG. **(A) t-SNE visualization for the distribution of MxG features** (after supervised dimension reduction) representing the 12 commands for a single session is shown for the user with best accuracy. Each dot represents a single trial, and each color represents a command. Note, the distinct clustering of individual commands in a user-dependent setting shows the significance of these features to be used for classification. **(B) Confusion matrix** for user-dependent classification accuracy, averaged and normalized across 9 users, is shown in a heatmap plot, with the accuracy of each command in the respective cell. Most misclassifications occurred between commands containing similar phonemes (e.g., ‘**i**ncrease volume’ vs ‘**de**crease volume’). **(C) Optimal sensor selection** using a forward selection strategy showed that about 7 optimal sensors were needed to achieve user-dependent accuracy above 80% (comparable to ExG+IMU performance in (Srivastava et al., 2024)). Even with a single sensor, average accuracy surpasses chance level (shown by a red line). Legends show the best participant performance. Dotted colored lines represent individual accuracy trends, and the solid black line shows the average. **(D) Accuracy across frequency bands:** [1–50], [50–100], [100–200], [200–300], and [1–300] Hz, all showed significantly higher than chance accuracy ([50–100 Hz: accuracy = 37.46%; one-sample *t*-test: *t*-stat = 9.76, *p* = 1.02 × 10^*−*5^; 100–200 Hz: accuracy = 35.44%; *t*-stat = 10.46, *p* = 6.06 × 10^*−*6^; 200–300 Hz: 24.38%; *t*-stat = 6.63, *p* = 1.65 × 10^*−*4^]), highlighting the importance of high-frequency MxG in SSR. Yet, 1–50 Hz suffices for classification ((1-way ANOVA: *F* = 1988.52, *p* = 7.48 × 10^*−*38^; post-hoc paired *t*-tests with Bonferroni correction: *t*-stat = 1.26, *p* = 0.25) vs 1–300 Hz).

Importantly, with sensors only around the ear the average accuracy was 74.87% (SD = 8.76%), and with sensors only on the face, it was 71.45% (SD = 7.86%) (Supplementary Figure S2-B, both higher than ExG alone (*∼* 70%). Note, the lower accuracy of face sensors could be due to a lower number of sensors at the face compared to around the ear (Supplementary Figure S2-A. All of these results were obtained using only 40 trials per command, indicating the practicality of the approach even with limited data. To better understand the impact of data quantity, we conducted an analysis of the data dependency and observed a clear trend: accuracy improved from approximately 50% with just 5 trials to about 80% with 20 trials and to about 86% with the full 40 trials (Supplementary Figure S3), demonstrating that further performance gains are likely achievable with larger datasets.

We also investigated the minimal number of sensors needed to maintain high accuracy. Using a forward selection strategy—starting from the best performing individual sensor and iteratively adding sensors that most improved classification—we found that only 7 sensors were sufficient to retain over 80% average accuracy (Figure 2C). Looking at the optimal sensor set for each user, we found that the sensors on the face and behind the ear had a higher frequency in the optimal set (Supplementary Figure S2-C). The ranking of all sensors across all users revealed a similar pattern (Supplementary Figure S2-D). Overall, this result suggests that effective silent speech recognition with magnetometers can be achieved with a significantly reduced number of sensors, contrasting with many existing physiological sensing approaches that often require dense electrode arrays or precise placement (Zhu et al., 2020), improving the practicality and cost-effectiveness of the system.

Additionally, we investigated how different frequency ranges contribute to SSR. While prior work on wearable SSR has primarily focused on frequencies below 50 Hz to capture subtle facial vibrations (Srivastava et al., 2024), we explored whether high-frequency muscle activity could also play a significant role. We compared performance across five distinct frequency bands — [1 *−* 50 Hz], [50 *−* 100 Hz], [100 *−* 200 Hz], [200 *−* 300 Hz], and [1 − 300 Hz]. Our results showed that MxG signals within the 1−50 Hz range alone were sufficient to achieve classification accuracies comparable to those obtained using the full broadband range [1−300 Hz] (Figure 2D), emphasizing the importance of accurately capturing muscle vibrations with magnetometers. Interestingly, signals from frequencies above 50 Hz also yielded classification accuracies significantly higher than chance (Figure 2D), highlighting that high-frequency muscle activity contains additional discriminative information. While low-frequency facial vibrations were adequate for this relatively simple command set, incorporating higher-frequency components could offer further improvements in SSR performance for more complex tasks.

### 2.2 User adaptation enables scalable SSR with headphone-MxG

We evaluated user-independent (training the model on data from a set of users and evaluating on a completely new user) performance with magnetometers in a headphone form factor, and also employed a curriculum learning strategy for user adaptation. Despite the increased variability introduced by cross-subject data, the features still formed 12 distinct clusters corresponding to the 12 silent commands in a t-SNE feature space after supervised dimension reduction (Figure 3A), demonstrating the generalizability of MxG representations.

**Figure 3:**
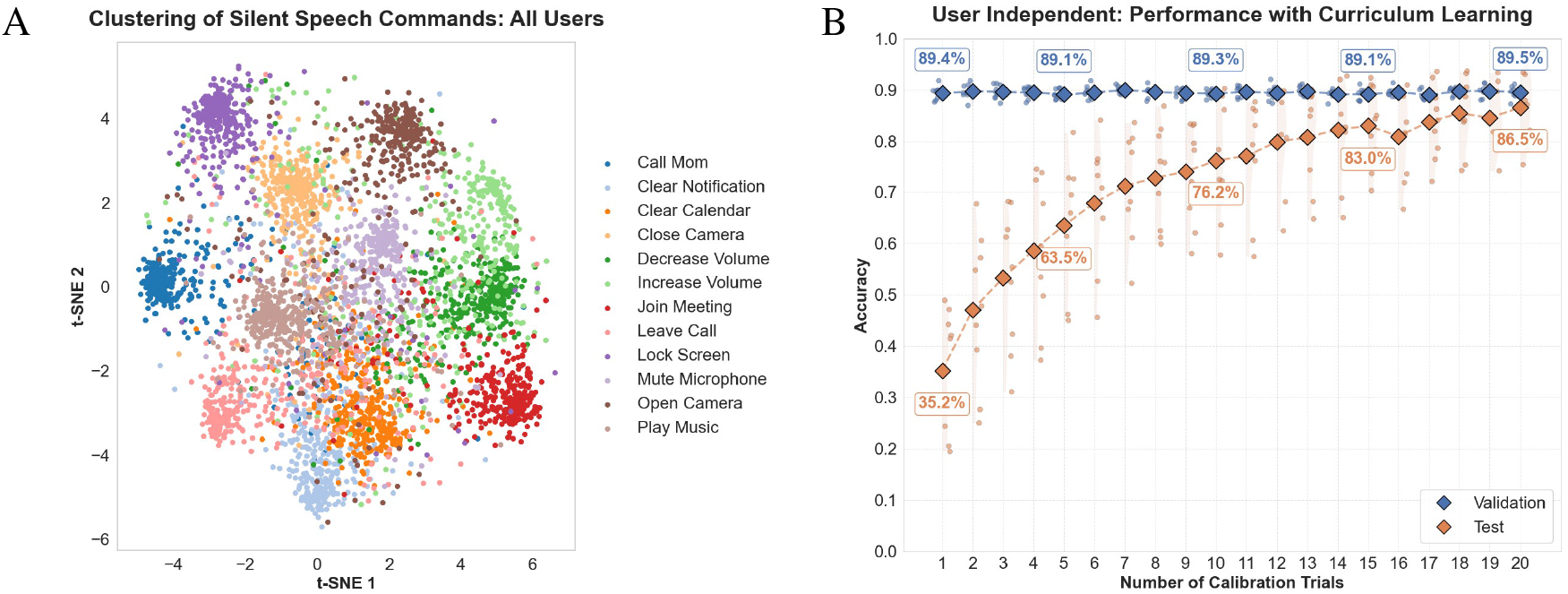
User-independent SSR performance with headphone-MxG. **(A) Visualization of MxG feature distribution across all participants**. t-SNE visualization after supervised dimension reduction of covariance features computed from the MxG signals of all 9 participants is shown. Each dot represents one trial, and each color represents a silent speech command. Note, the emergence of 12 distinct clusters representing the 12 classes, indicating the generalizability potential of MxG features. **(B) User-independent SSR accuracy trend with increasing user adaptation** Validation and test accuracy trend for user-independent SSR with increasing number of user adaptation trials using curriculum learning is shown. The blue curve shows the validation accuracy, and the orange curve shows test accuracy for each number of calibration trials. Note, test accuracies were computed from the held-out user data, after taking out the calibration trials. Dots represent accuracy distribution of all users. Note that, with about 12 calibration trials, the test accuracy reaches above 80%, closer to user-dependent performance.

Given the small sample size (N=9), training a user-independent deep learning model without any user adaptation yielded an accuracy of 25.9%, much lower than the user-dependent performance (86%), though still significantly above chance. This gap highlights the presence of variability in the captured MxG signals, likely stemming from differences in sensor placement due to variations in head size, which remains a critical challenge for future consumer-facing SSIs. However, as seen in many largescale machine learning systems, it is expected that with greater amounts of similar training data, the model will eventually learn to generalize across such variability (Hestness et al., 2017; Rosenfeld, 2021). To illustrate such potential for improvement, we applied curriculum learning-based user adaptation, retraining the model with increasing numbers of subject-specific calibration trials added to the training set and testing with remaining held-out user trials. Performance improved markedly, reaching 63.5% test accuracy with just 5 calibration trials, 76.2% with 10 trials, 83.0% with 15 trials, and 86.5% with 20 trials (Figure 3B). This trend was also observed in the learning curves as the reduction of the gap between training-, validation-, and test-accuracy with increased user adaptation (Supplementary Figure S4). Different types of curriculum learning strategy yielded similar results (Supplementary Figure S6). Crucially, high-frequency MMG achieved much higher user-independent performance compared to user-dependent settings, highlighting MMG’s reduced cross-subject variability characteristics at high frequency (Supplementary Figure S7). These results demonstrate that even minimal user-specific calibration can substantially boost performance, making magnetometer-based SSIs scalable and highly promising for real-world deployment.

### 2.3 Magnetometers in a necklace form factor can decode both overt and silent speech in ambient conditions

Beyond facial vibrations, overt speech also generates bone-borne vibrations originating from the vocal cords, which propagate through skeletal and muscular pathways, particularly along the neck and jaw (Marur et al., 2014; McBride et al., 2008). The necklace form factor is particularly well-suited to capture these signals, offering an optimal anatomical location for sensing both neuromuscular activity and bone-conducted mechanical vibrations during overt speech.

Motivated by the physiological insight of prior overt EMG works leveraging neck muscles in a wearable form factor (Wu et al., 2024), we explored whether magnetometers embedded in a necklace form factor could similarly capture the rich articulatory dynamics present during overt speech and further, to what extent this configuration could support effective silent speech recognition despite the absence of large vocal fold-driven vibrations. We adapted the stimuli from (Wu et al., 2024) (Figure 4A) for both overt and silent speech classification.

**Figure 4:**
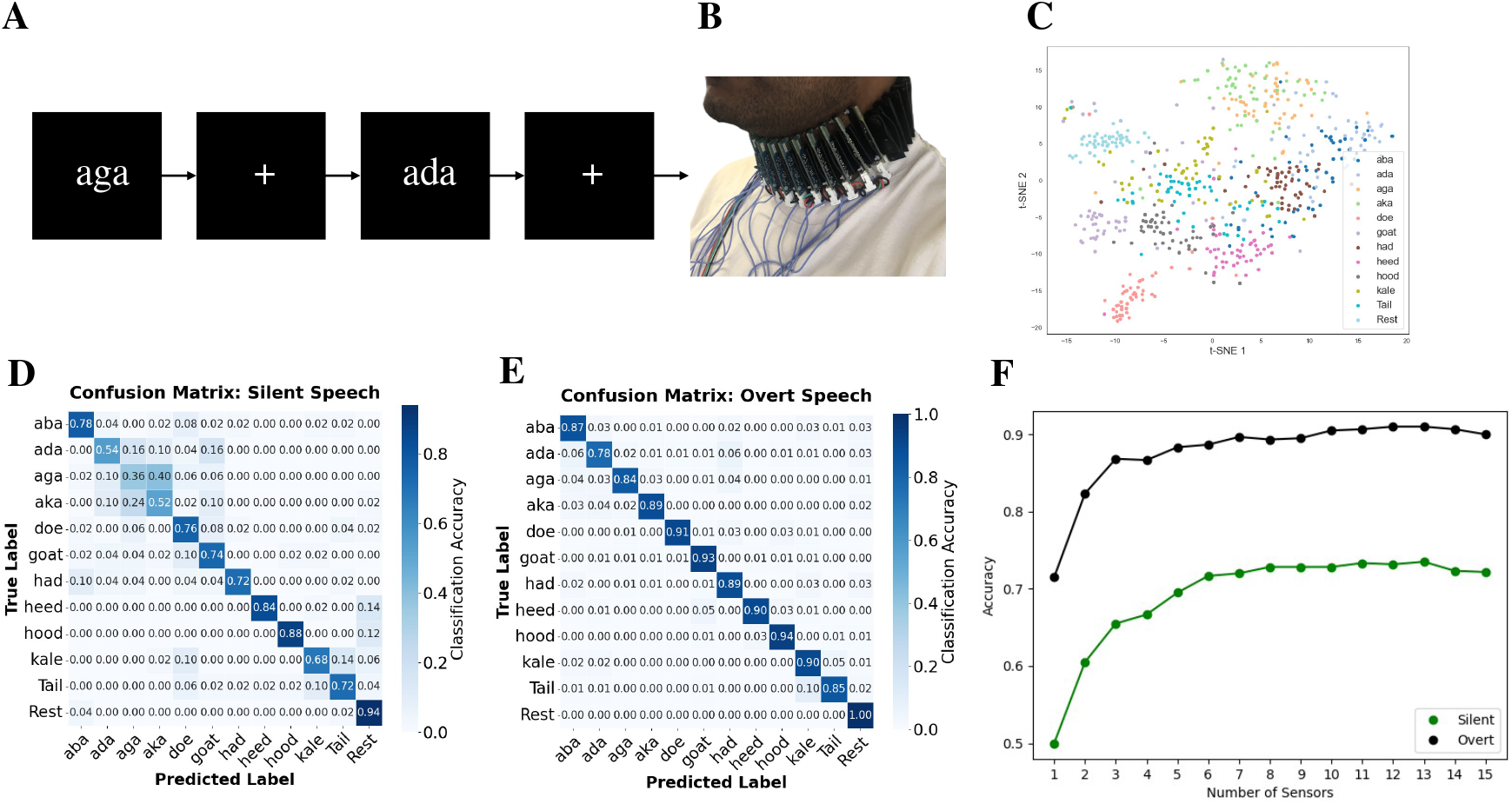
Overt and Silent speech decoding with Necklace-MxG. **(A) The Similar Words’ task for both silent and overt speech with necklace form factor**. The task was designed similarly to the EMG-Necklace study (Wu et al., 2024), where participants (N=2) overtly and silently articulated 11 similar words: ‘heed’, ‘had’, ‘hood’, ‘tail’, ‘kale’, ‘doe’, ‘goat’, ‘aba’, ‘ada’, ‘aga’, and ‘aka’. Each word was displayed on a computer screen for 1.5 s followed by 1.5 s of neutral period (+). Each participant performed 2 sessions, one for silent speech and one for overt speech, with 50 trials per session. **(B) Equipment for data collection**. 16 Necklace worn by one of the authors for demonstration. Aichi DJ single-axis magnetometers were placed on a custom 3D printed flexible platform to be wrapped around the front side of the neck with their sensitive axis radial to the neck. The data was collected in ambient conditions in a typical office setting. **(C) Visualization of the features**. User-dependent t-SNE visualization of the covariance features extracted from the MxG signals showed distinct clustering of different classes. Each color represents a word stimulus class, and each dot represents a trial. **(D) Confusion matrix for silent speech classification**. Using a linear classifier and 8-fold cross-validation strategy, user-dependent classification of the 11 words resulted in 71% accuracy for silent speech as shown in the confusion matrix, averaged across participants. Note the major confusion between ‘ada’, ‘aga’, and ‘aka’ that differed in only a plosive place of articulation, which is difficult to capture at the neck. **(E) Confusion matrix for overt speech classification**. Similar user-dependent classification strategy for overt speech resulted in 90% classification accuracy for the 11 similar words, significantly higher than silent speech and on par with the overt EMG Necklace study (Wu et al., 2024). **(F) Optimal sensor selection accuracy trend**. A forward selection strategy resulted in about 7 sensors, after which both overt and silent speech recognition performance plateaued.

A major challenge in the deployment of magnetometers for wearables has historically been their sensitivity to the magnetic field of the Earth, which required operation within magnetically shielded environments, as in the case of our earlier OPM experiments. However, continued progress in increasing dynamic range has resulted in magnetometers that can operate in ambient conditions. We developed a magnetometer-based necklace system (Figure 4B) using Aichi DJ sensors (Aichi Steel Corporation), which are compact and capable of measuring subtle magnetic fluctuations generated during articulatory movement in ambient conditions.

To visualize the discriminative potential of the MxG signals, we extracted time-domain covariance features from the magnetometer data and applied t-SNE projection. As shown in Figure 4C, t-SNE visualization revealed clear clustering of trials corresponding to different spoken words. Each color represents a different word class, and each dot corresponds to a single trial. Despite the small sample size and ambient recording conditions, the clusters were distinct, indicating that the magnetometers effectively captured word-specific MxG patterns from the neck during overt articulation.

We evaluated word classification performance using a linear classifier with an 8-fold cross-validation strategy. Classification of overt speech trials resulted in an accuracy of 90%, aligning closely with prior results from the EMG-Necklace study (Wu et al., 2024), whereas for silent speech the accuracy was about 71%. Analysis of the silent speech confusion matrix (Figure 4D) indicated that the majority of classification errors occurred among the words ‘ada’, ‘aga’, and ‘aka’, which share highly similar articulatory features distinguished only by subtle differences in place of articulation. These results suggest that while silent articulation imposes challenges for neck-mounted sensing, particularly for phonetically similar sounds, magnetometers can still achieve meaningful SSR performance. But, for overt speech (Figure 4E), misclassifications were minimal, demonstrating that overt speech produces stronger and more distinct magnetic signatures at the neck compared to silent articulation. These findings validate the use of neck-mounted magnetometers as a promising sensing modality for both silent and overt speech recognition.

In addition, to investigate the optimal number of sensors required, we used the same forward selection strategy. For both silent and overt speech conditions, the accuracy steadily improved as more sensors were incorporated. However, after the inclusion of approximately 7 sensors, the accuracy plateaued, indicating diminishing returns with more sensors. This suggests that a relatively small subset of sensors is probably sufficient to capture the minimal information necessary for closed-vocabulary speech recognition, such as this task. Importantly, identifying this minimal set of sensors not only supports the development of lightweight and cost-effective wearables but also reduces computational complexity and energy demands for real-time silent speech decoding.

## 3 Discussion

In this study, we introduced magnetometer-based sensing as a powerful and scalable alternative for wearable SSR. By embedding OPMs into a headphone form factor, we demonstrated that MxG signals can achieve user-dependent classification accuracy of about 86% across 12 distinct silent speech commands—substantially outperforming previously reported ExG+IMU systems. Through curriculum learning-based user adaptation, we reduced training demands to as little as 5 minutes and calibration demands to as few as 12 trials (18 seconds) per command while maintaining high accuracy. Our forward sensor selection analyses further revealed that robust classification could be achieved with as few as 7 optimally placed sensors, highlighting the feasibility of cost-effective and wearable system designs. Moreover, we established that both overt and silent speech can be reliably decoded in ambient environments using a flexible neck-worn magnetometer array, achieving 71% (silent) and 90% (overt) classification accuracy for phonetically similar words. Collectively, these results position magnetic sensing as a promising foundation for the next generation of consumer-centric silent speech interfaces.

### 3.1 Magnetometers as the preferred alternative modality for wearable SSR

Despite the range of sensing modalities explored for silent speech interfaces (SSIs)—including surface electromyography (sEMG) (Meltzner et al., 2018; Song et al., 2023; Wu et al., 2024; Zhu et al., 2020), inertial measurement units (IMUs) (Kwon et al., 2023), and audiovisual techniques (K. Sun et al., 2018)—SSIs remain limited by several practical and technical challenges that hinder their wearability and real-world applicability. Magnetometers, by contrast, offer a compelling alternative: they enable the non-invasive detection of minute biomagnetic signals generated by articulatory muscle activity without requiring direct contact with the skin (Cohen & Givler, 1972; Yun, Gonzalez, et al., 2024), thereby improving both wearability and user convenience. In our work, embedding magnetometers into a headphone form factor enabled 86% user-dependent classification accuracies (Figure 2), significantly surpassing prior state-of-the-art wearable SSI systems, including those combining ExG and IMU modalities (Srivastava et al., 2024). This performance suggests that magnetometers effectively capture fine-grained articulatory dynamics, rivaling if not exceeding, traditional biosensors such as sEMG.

Our finding of classification performance exceeding prior benchmarks using fewer and more flexibly placed sensors (Figure 2C, Supplementary Figure S2) suggests that magnetic sensing can achieve high-resolution decoding with reduced hardware complexity, enabling designs that are not only less obtrusive but also better suited for wearable deployment. The strong performance of the ear-only configuration in this study along with the previous studies showing reliable SSR with sEMG sensors around the ear (Jin et al., 2022; Srivastava et al., 2024; X. Sun et al., 2024) demonstrate the anatomical richness of the peri-auricular region for capturing articulatory dynamics and supports the development of SSI systems embedded into common wearable form factors like headphones or earbuds. Even in the face-only configuration, despite having fewer sensors, magnetometers outperformed ExG-only baselines, suggesting that magnetic signals capture more robust or discriminative features even with suboptimal coverage.

This highlights a critical advantage of magnetometry: the ability to sense facial muscle activity without requiring direct skin contact. This is in contrast to sEMG-based systems, which often require dense electrode arrays for similar tasks and perform poorly when integrated into hair-covered regions like the scalp or jawline (Zhu et al., 2020). Magnetometers, with their ability to sense through hair or even clothing and without requiring conductive gel or direct skin contact (Jonna & Rao, 2023; Yun, Gonzalez, et al., 2024), unlock form factors—such as headphones, earbuds, and smart glasses—that are simply not viable with traditional ExG-based approaches. Thus, the success of the headphone form factor highlights the feasibility of embedding magnetic sensing into familiar consumer electronics with only minor hardware modifications, supporting discreet and accessible SSI solutions. More broadly, these results mark a meaningful shift in the wearable SSR design paradigm from clinical-grade, contact-dependent systems to practical, user-friendly form factors capable of supporting everyday silent communication.

Beyond raw performance, our findings are meaningful in light of the broader challenges facing the field. Many prior studies using high-density EMG or brain-based methods (e.g., EEG) have been limited to small vocabulary or highly constrained tasks (Ghane et al., 2015; Min et al., 2016; Zhang et al., 2025; Zhu et al., 2020). Brain recordings, while promising for open-vocabulary decoding, require invasive hardware (Meng et al., 2023) or suffer from poor signal-to-noise ratios in noninvasive setups (Min et al., 2016; Pei et al., 2012). High-density EMG, although capable of capturing high-resolution neuromuscular activity, is difficult to integrate into wearable solutions (He et al., 2023; Milosevic et al., 2017). Our results suggest that magnetometer-based wearable systems can capture rich articulatory cues, opening a path toward larger vocabulary spaces while preserving the wearability and practicality required for consumer adoption.

### 3.2 Magnetic sensing enables scalable, personalized silent speech interfaces

Scalability and ease of personalization remain the central challenges in the development of practical SSIs using electrophysiological sensors (Deng et al., 2014; Pandey et al., 2021). While our initial user-independent performance with only 5 minutes of training data was modest (26% accuracy), we demonstrated substantial gains to over 63%*−* 83% with only 5 *−*15 calibration trials through user adaptation (Figure 3). This rapid adaptation trend reflects a fundamental advantage of magnetic sensing: MxG signals offer higher signal-to-noise ratios (Cohen & Givler, 1972), less dependence on skin conditions (Yun, Gonzalez, et al., 2024), and more consistent anatomical targeting compared to traditional surface electrodes (Elzenheimer et al., 2020; Yun, Csaky, et al., 2024), than ExG modalities like EMG, which are often confounded by skin impedance, contact variability, and facial hair. Moreover, because magnetometers do not require direct skin contact and operate in three dimensions, they can be placed with greater spatial density—even in compact regions—enabling higher-dimensional signal capture and, in turn, more expressive and accurate SSR.

Prior work on gesture recognition has shown that magnetometers allow user-independent gesture recognition that outperforms traditional EMG systems in both accuracy and generalizability (Yun, Csaky, et al., 2024). Similarly, for SSI applications, we demonstrated user-independent SSR performance close to user-dependent performance, with as little as 5 minutes of training data and 18 s of calibration data per silent command. These results suggest that MxG could fundamentally shift the scalability paradigm from training thousands of hours of data toward minimal training and adaptation requirements. Even limited calibration data can be highly informative when paired with stable and information-rich MxG features, which are a critical factor for real-world deployment where users may tolerate only brief calibration periods. By leveraging efficient user adaptation and new strategies to further boost generalization—such as subject-agnostic pretraining (Zhou et al., 2024), domain adaptation (Farahani et al., 2021), federated learning (Mammen, 2021), few-shot fine-tuning (Liu et al., 2022), and combining with audiovisual techniques—future magnetometer enabled SSIs could offer greater scalability, greatly expanding accessibility to broader populations. Such optimizations are essential for achieving the scale and simplicity needed for widespread consumer adoption of MxG-enabled SSI technologies.

### 3.3 Silent speech recognition with magnetometers can be achieved in unshielded, real-world conditions

Previous HCI studies involving magnetic sensing have been constrained to magnetically shielded environments (Greco et al., 2023; Yun, Csaky, et al., 2024), limiting their translational potential. In contrast, our findings demonstrate that silent and overt speech can be reliably decoded from the neck using magnetometers under fully ambient conditions. This was made possible by leveraging advancements in magnetometer design, particularly sensors with enhanced dynamic range, combined with a neck-mounted form factor optimized for articulatory signal capture. Despite using non-specialized, off-the-shelf sensors, we achieved 71% accuracy for silent speech and 90% for overt speech across 11 phonetically similar words, rivaling state-of-the-art wearable EMG systems like the EMG Necklace (Wu et al., 2024).

Future improvements in sensor density, sensitivity, and dynamic range, as well as multimodal fusion strategies (e.g., headphone + necklace form factor) could further enhance spatio-temporal resolution of muscle sensing. As novel magnetometers such as acoustically driven ferromagnetic resonance (ADFMR) sensors (Labanowski, 2017) continue to improve by becoming smaller, more sensitive, lower power, and better suited for body integration, the path toward consumer SSIs become increasingly feasible. Findings in this study provide the first proof of concept that magnetometers can reliably support SSR under everyday ambient conditions, a critical milestone for realizing practical, wearable SSR solutions using magnetic sensors.

### 3.4 Future directions: ubiquitous wearable magnetic sensors and practical applications

As silent speech recognition (SSR) technologies mature, their real-world relevance becomes increasingly clear across a range of practical domains. From assistive communication for individuals with speech impairments to hands-free control in AR/VR environments, and secure, noiseless exchange in covert operations, the potential applications are diverse and compelling. These use cases highlight the need for SSR systems that go beyond accuracy to prioritize wearability, discretion, and everyday usability, thus, positioning wearables with magnetic sensors as a particularly promising path for future innovation.

A clear next step for practical deployment is to adapt magnetic sensing to wearable form factors in real-world, ambient environments. Achieving this will require optimizing sensor placement around the ear to maximize articulatory signal capture while minimizing interference from environmental magnetic noise. Advances in chip-scale magnetometer technologies such as ADFMR sensors (Labanowski, 2017) will be critical to enabling lightweight, low-power, and fully wearable designs that can be seamlessly integrated into everyday accessories such as earbuds and glasses. Signal processing pipelines that can adapt to ambient magnetic conditions will also be crucial.

In parallel, moving toward open-vocabulary SSR presents a distinct set of challenges. This shift will necessitate richer training datasets, improved phoneme-level decoding models, and potentially hybrid sensing strategies that integrate magnetic sensing with complementary modalities, such as inertial sensing for enhanced robustness, adaptability, and generalization. Techniques such as domain adaptation, meta-learning, and continual learning could greatly minimize calibration requirements by addressing inter-session and inter-user variability. By advancing the scalability, robustness, and usability of magnetometer-based wearable SSIs, this work moves the field closer to practical, real-world deployment—opening the door to a future where wearable silent speech recognition becomes an integral part of everyday communication technologies. Magnetometer-based SSIs have the potential not only to augment human communication, but also to fundamentally reshape how we interact with technology, each other, and the world around us.

## 4 Methods

### 4.1 Data Collection

#### 4.1.1 Magnetically shielded room (MSR)

All experiments were conducted inside a magnetically shielded room (Magnetic Shield Corporation, MuROOM) with internal dimensions of 1.3 × 1.3 × 2 cubic meters. The MSR provided approximately 25, 000 fold attenuation of residual magnetic fields at DC and up to 8, 000 fold attenuation at AC, minimizing contamination from ambient noise sources. Before each recording session, the room was degaussed by applying a decreasing sinusoidal field through embedded coils, effectively removing any wall magnetization and maximizing ambient field rejection. A projector mounted outside the MSR displayed visual stimuli on a screen inside the room through an opening in the wall.

#### 4.1.2 Sensors

We used two-axis QZFM Gen-2 optically pumped magnetometers (OPMs; QuSpin, Inc.) and single-axis Aichi DJ sensors (Aichi Steel Corporation) for MxG measurements.

##### Headphone form factor

The OPM sensors were housed in a 3D-printed headphone structure containing 10 sensors on each side, with 7 sensors around the ear and 3 on the face (Supplementary Figure S2-A). Each OPM was sensitive along two orthogonal axes, positioned so that one axis was tangential to the face surface and the other radial to the face. Recordings were made from both axes of each sensor. Thus, data collection involved 20 OPM sensors (10 per side), with two axes per sensor, totaling 40 channels. The headphone design was adapted from open-source designs available at homebrewheadphones.com and modified in Fusion 360 to incorporate custom housings for the OPMs. Sensors were secured in place using nylon M2 screws.

##### Necklace form factor

16 Aichi DJ sensors were embedded into a custom-made flexible 3D-printed holder, with each sensor oriented such that its sensitive axis was radial to the neck. The band of sensors covered the entire front half of the neck, roughly extending from one ear to the other. Additional custom hardware was designed to ensure clock synchrony between the sensors to limit crosstalk, and to route all analog output signals to an ADC. An analog high-pass filter with 20 Hz cutoff was used to prevent sensor saturation.

#### 4.1.3 Data Acquisition

All sensors output analog signals were digitized using 16- or 24-bit NI-DAQ ADC modules (either NI-9202 or NI-9205 modules in a cDAQ chassis, National Instruments) at a sampling rate of 2 kHz per channel and collected with custom Python data acquisition code.

#### 4.1.4 Tasks

Data collection involved two tasks: (1) ‘Commands’ task with OPMs embedded in a headphone form factor for silent speech only, and (2) ‘Similar Words’ task with Aichi magnetometers embedded in a necklace form factor for both silent and overt speech in ambient.

**Table 1:**
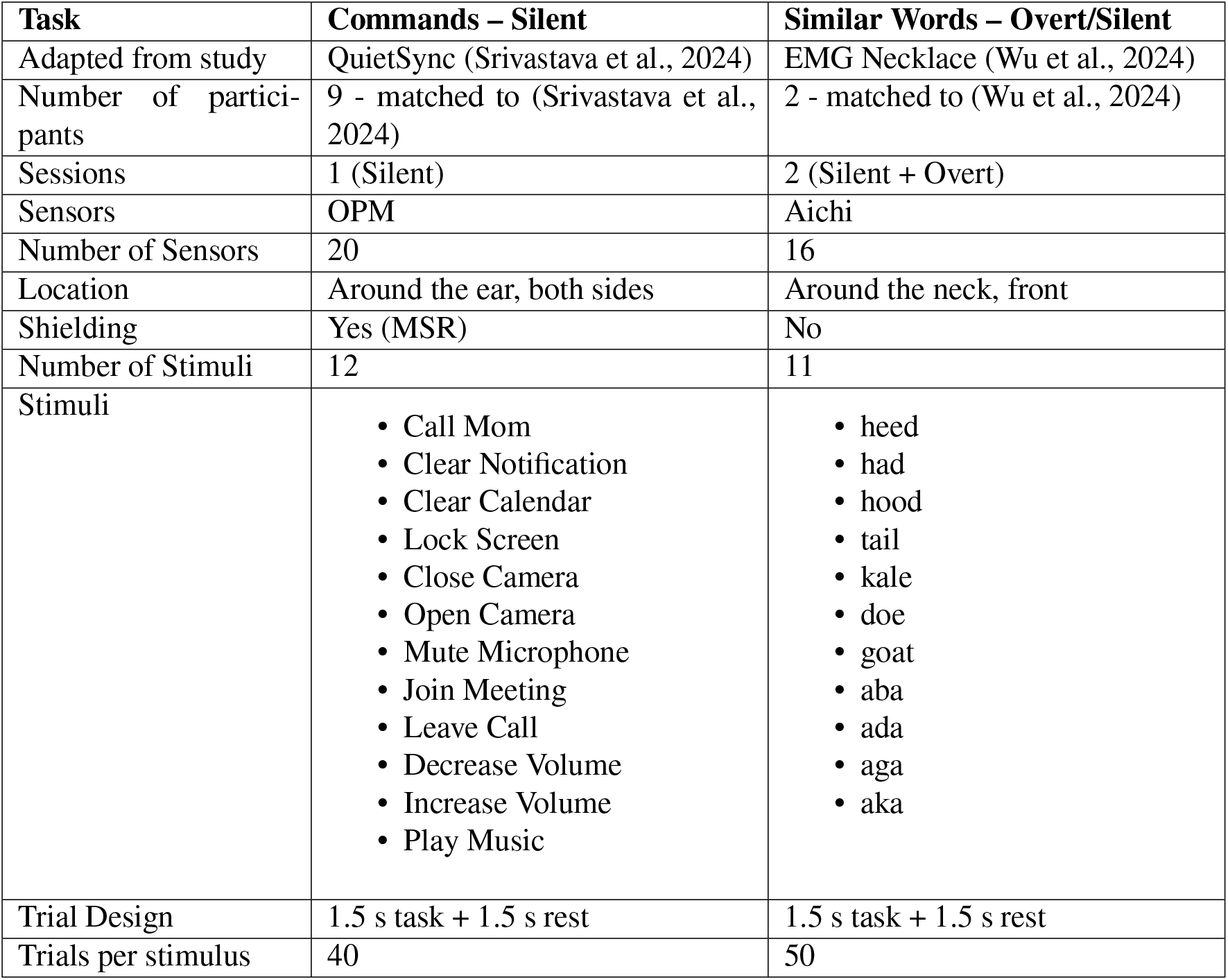
Task details.

##### Commands task

We used the same set of stimuli as in the QuietSync study with ExG + IMU (Srivastava et al., 2024). Participants silently articulated 12 commonly used speech interaction commands: *Call Mom, Clear Notification, Clear Calendar, Lock Screen, Close Camera, Open Camera, Mute Microphone, Join Meeting, Leave Call, Decrease Volume, Increase Volume*, and *Play Music*. This task was conducted inside the magnetically shielded room (MSR) using OPM sensors in a headphone form factor. Before starting the task, the MSR was demagnetized (degaussed), and the OPMs were calibrated while a participant was in the room. The stimuli were projected onto a screen via an external projector placed outside the MSR. Each command was repeated for 40 trials in a pseudo-randomized order, with each trial consisting of a 1.5-second task period followed by a 1.5-second rest period.

##### Similar Words task

The same stimuli as in the EMG Necklace study (Wu et al., 2024) were used: *heed, had, hood, tail, kale, doe, goat, aba, ada, aga*, and *aka*. These utterances have been designed to target specific aspects of speech variation. They include *minimal pairs* differing only in vowel identity (e.g., [hid], [hæd], [hUd]); *Near-minimal pairs* differing in the place of articulation of a plosive (e.g., [aba], [ada], [aga]; [doU], [goUt]; [DEIl], [kEIl]); and pairs that contrast voicing and aspiration in plosives (e.g., [aga] vs. [akha]). These structured contrasts enable the isolation of individual speech features, allowing for more precise evaluation of articulatory and acoustic differences. The “Similar Words” task was performed in ambient conditions (i.e., without magnetic shielding), using Aichi DJ sensors. The stimuli were displayed on a monitor, and participants performed two sessions: one *overt*, where they read the stimuli aloud, and one *silent*, where they mouthed the words without vocalization. For this task, data were collected in a manner similar to the previous command task: the stimulus was displayed for 1.5 s, followed by 1.5 s of rest (neutral).

#### 4.1.5 Participants

Nine participants (7 males and 2 females; 25 *−* 64 years in age; 2 non-native English speakers; 2 with facial hair) participated in the silent ‘Commands’ task. 2 participants (Males, 29 and 33 years of age; 1 non-native English speaker) participated in the ‘Similar Words’ task. All participants are employees of Sonera. Prior to the start of data collection, the participants were informed that they could exit the study at any time and for any reason. It was ensured that no participants had any ferrous material on them including any jewelry or metallic dental work such as permanent retainers or braces in their mouth. All participants were consented to and were given instructions before collecting data, and were compensated for their time. To get familiarized with the task, participants practiced the task for a few trials prior to the start of the experiment. They were seated in a non-ferrous chair inside the MSR for the ‘Commands’ task. Personally identifiable information (PII), including demographic information (name, email address, phone number) and biometric information (physiological signals, age, and additional information) was housed and recorded separately following approved confidentiality protocols.

### 4.2 Data analysis

#### 4.2.1 Preprocessing

All data were recorded at a sampling rate of 2 kHz. To eliminate line noise, we applied a 2^*nd*^ order IIR notch filter targeting 60 Hz harmonics (Q-factor = 30). For analyses requiring broader spectral content, signals were bandpass filtered between 1 *−* 300 Hz using a 5^*th*^ order Butterworth filter.

#### 4.2.2 Feature extraction

For each trial, we extracted covariance-based features to capture spatial relationships across channels. Preprocessed MxG signals were organized as a two-dimensional array: channels (40 for OPMs; 16 for Aichis) × time samples (e.g., 2000; equivalent to 1 s (after correcting for reaction time) at 2 kHz). We computed the covariance matrix across channels, capturing pairwise linear dependencies indicative of coordinated activity. To reduce dimensionality and redundancy, we retained and vectorized only the upper triangular portion of the covariance matrix, producing a compact yet informative feature vector per trial. This representation proved effective for classification tasks by encoding cross-channel interactions.

#### 4.2.3 Visualization

To qualitatively assess the discriminability of the extracted features across silent speech commands, we employed t-SNE visualization (Van der Maaten & Hinton, 2008). Covariance features from all participants were compiled into a matrix alongside their class labels. Linear Discriminant Analysis (LDA) was used to project the high-dimensional feature space into a maximally discriminative subspace (dimensionality = number of classes - 1). The resulting features were normalized via Min-Max scaling, then embedded into two dimensions using t-SNE (with PCA initialization and correlation distance metric). This configuration was selected to preserve both local and global structure while emphasizing inter-class separation. The 2D embeddings were visualized as a scatter plot, with each point representing a trial colored by its associated command label. The resulting clustering patterns suggested that the covariance features capture discriminative information relevant to speech command classification.

#### 4.2.4 User-dependent classification

To assess the classification performance of silent speech decoding in a user-dependent setting, we implemented a comprehensive machine learning pipeline incorporating all available supervised classifiers from the scikit-learn library (Supplementary Figure S1). For each participant, models were trained and evaluated exclusively on that individual’s MxG data, simulating a scenario in which systems are calibrated and used on a per-user basis. A subset of classifiers was excluded from analysis based on their incompatibility with the task (e.g., multi-class limitations, assumptions of categorical features, or excessive computational requirements). For models supporting the max-iter parameter, this value was set to 1000 to ensure adequate convergence.

The features were standardized using z-score normalization. Classification performance was evaluated using an 8-fold stratified cross-validation scheme. In each fold, the training data were used to fit the model, and predictions were made on the held-out fold. Accuracy was computed for each fold, and the mean and standard deviation across folds were recorded. Confusion matrices were also computed per fold and aggregated to produce a subject-level summary of classification performance for each classifier. This systematic framework enabled a comprehensive comparison of classifier effectiveness across both individual users and classifier types.

The classifier resulting in the best average accuracy across users was RidgeClassifier, which was selected for further analysis with hyperparameter tuning with nested cross-validation. To optimize the RidgeClassifier for user-dependent classification, we implemented a two-stage hyperparameter tuning process. The following table outlines the steps taken during each stage of optimization.

This two-tiered approach allowed for efficient narrowing of the search space and fine-tuning of model behavior to capture subject-specific characteristics in the silent speech decoding task.

##### Sensor Selection

To identify optimal sensor subsets for user-dependent silent speech decoding, we implemented a progressive sensor selection procedure using RidgeClassifier and stratified 8-fold cross-validation. Each of the 20 physical sensors included two channels, totaling 40 input channels per participant. For each subject, classification performance was iteratively evaluated by incrementally adding one sensor (i.e., two channels) at a time. At each step, all possible additions of a new sensor to the previously selected set were evaluated based on mean classification accuracy. For each candidate subset, features were extracted from the selected channels using covariance-based features over a task-specific temporal window determined by the highest performing onset per subject. Features were standardized using z-score normalization, and model performance was assessed via accuracy and confusion matrix metrics across folds.

**Table 2:** Hyperparameter optimization for Ridge Classifier.

| Stage | Component | Parameter | Values / Description |
| --- | --- | --- | --- |
| Preprocessing | Feature Selection | VarianceThreshold | Remove low-variance features |
|  | Standardization | StandardScaler | Z-score normalization |
|  | Univariate Selection | SelectKBest (score_func = ANOVA F-score) | Select top-k features based on univariate test |
| Stage 1 | Feature Selection | feature_select_k | {0.2, 0.5, 0.8, 'all'} (proportion of features retained) |
| | Regularization Strength | alpha | Log-spaced values from $10^{-6}$ to $10^6$ |
|  | Solver | solver | {'auto', 'svd', 'lsqr', 'saga', 'sparse_cg'} |
| | Convergence Criteria | tol | $10^{-4}$ (fixed) |
| Stage 2 | Iteration Limit | max_iter | 1000 (fixed) |
|  | Positive Coefficients | positive | {True, False} |
|  | Fit Intercept | fit_intercept | {True, False} |
| | Fine Tuning of Tolerance | tol | $\{10^{-5}, 10^{-4}, 10^{-3}, 10^{-2}\}$ |

The best-performing sensor combination at each iteration was retained, allowing us to construct a subject-specific ranked list of sensors based on incremental contributions to classification performance. This process enabled efficient identification of compact sensor subsets that maximized accuracy with minimal channel count. Note, sensor selection was conducted independently within each training fold, and classification performance was evaluated on the corresponding held-out fold. This yielded an unbiased estimate of performance for each sensor count. Accuracy trends across sensor counts and corresponding sensor combinations were recorded for each subject, enabling both performance benchmarking and potential hardware reduction for practical deployments.

#### 4.2.5 User-independent Classification

To develop a robust silent speech recognition (SSR) model capable of generalizing across different users, we designed a deep learning framework that integrates depthwise separable convolutions (Kaiser et al., 2017), squeeze-excitation (SE) blocks (Hu et al., 2018), multi-scale attention mechanisms (Chen & Shi, 2021), and curriculum learning-driven domain adaptation (Bengio et al., 2009; Dash et al., 2023). The primary objective of the model is to learn hierarchical, subject-invariant features from multichannel MxG signals while maintaining high classification accuracy in a user-independent setting. Note that, first we attempted user-independent classification with linear methods, similar to the user-dependent strategy, but the linear models were expectedly unable to capture the user variability with such limited data (Supplementary Figure S5).

Preprocessed MxG signals at 1 *−* 300 Hz range downsampled to 1000 Hz sampling frequency were used for user-independent analysis. For comparison with different frequency ranges (Supplementary Figure S7), the MxG signals were bandpass filtered to respective cutoffs and downsampled to at least 4× range of high cutoff using polyphase resampling to reduce computational load while preserving signal fidelity. For example, for analysis with 1 *−* 50 Hz, MxG signals were downsampled to 200 Hz. Before training, the signals were standardized channel-wise to zero mean and unit variance.

The deep learning model was structured as follows:

##### Input Block

The downsampled signals are passed through an initial convolutional block. This consists of a 1D convolutional layer with 64 output channels, a kernel size of 15, and padding of 7 to capture local temporal dependencies. A Batch Normalization layer stabilizes activations, followed by a Gaussian Error Linear Unit (GELU) activation. This is followed by average pooling with a pool size of 4 to reduce temporal resolution. A 1 × 1 convolution for channel remapping and a second batch normalization layer are applied, and a residual connection is introduced by summing the input and the transformed feature map, improving gradient flow and model stability.

##### Depthwise Separable Convolution Blocks

Two sequential depthwise separable convolution blocks are used to progressively extract spatial and temporal features with high computational efficiency. Each block factorizes standard convolution into:

- **Depthwise convolution**: Applies a single convolutional filter per input channel.
- **Pointwise convolution**: Combines the outputs via 1 × 1 convolutions.

The first block uses a kernel size of 15 and is followed by max pooling (pool size 2) for further temporal downsampling. The second block uses a kernel size of 9 and includes adaptive average pooling, which compresses feature maps into a fixed temporal dimension of 25 steps, enabling consistent downstream processing across variable-length input sequences.

##### Squeeze-Excitation (SE) Blocks

To enhance the network’s capacity to model inter-channel dependencies, each depthwise separable block is paired with a Squeeze-and-Excitation (SE) block. SE blocks operate in two stages:

- Squeeze: Global average pooling collapses each channel’s temporal data into a scalar representing its overall importance.
- Excitation: A lightweight two-layer fully connected network (with a bottleneck reduction ratio of 8) predicts channel-wise weights, which are then used to recalibrate the feature maps via channel-wise multiplication.

This mechanism selectively emphasizes the most informative features and suppresses noise-prone channels, contributing to improved inter-subject generalization.

##### Multi-scale Attention

Following the SE-enhanced feature maps, a multi-scale attention module was applied. This allowed the model to dynamically focus on salient temporal segments at varying time scales. The attention mechanism operates on the output tensor (after permutation to Batch Size (B), Sequnce Length (L), Feature Size (C)), performing linear projections to obtain query, key, and value matrices, and calculates attention weights via scaled dot-product attention. These weights are used to compute a weighted sum of the values, and the result is mean-pooled across the temporal dimension to yield a condensed 256-dimensional representation per sample. This mechanism is crucial for modeling both short-term modulations and long-term dependencies in temporal patterns.

##### Classifier Block with Spectral Normalization

Finally, the classifier block translates the learned features into final predictions. It consists of a fully connected layer with 512 output units, followed by spectral normalization on the weight matrices to prevent overfitting and ensure stable learning dynamics. This is then followed by layer normalization, which normalizes the activations across feature maps, stabilizing training. A GELU activation function introduces non-linearity, and dropout with a rate of 0.5 is used to prevent overfitting by randomly deactivating neurons during training. A final fully connected layer (512 to the number of classes) produces the model’s output, which is implicitly passed through a softmax activation (by the loss function) to generate class probabilities.

##### Loss Function and Regularization

To optimize training, we used Label Smoothing Cross-Entropy loss, with a smoothing factor of 0.1. Label smoothing regularizes the model by preventing it from becoming overconfident in its predictions, making it more robust. Regularization techniques employed include Dropout (with a rate of 0.5 in the classifier block) and Spectral Normalization (applied to the first linear layer in the classifier).

##### Curriculum Learning

Curriculum learning is integrated into the training process to improve the model’s generalization performance. The fundamental idea is to gradually introduce more difficult examples as the model becomes more proficient. This staged approach helps the model build a solid foundation of generalizable features before being exposed to more challenging examples. The curriculum adjusts the proportion of samples included in each epoch’s training data. This “difficulty factor” dynamically adapts based on the current epoch progress (ramping up from 0.2 to 0.9 of the full dataset) and is further modulated by the model’s recent validation performance and the class balance observed in the validation set. Specifically, if the model struggles (low recent accuracy and negative trend), the difficulty factor is reduced (made easier), and if it performs well (high recent accuracy and positive trend), it is increased (made harder). Samples within the selected subset for each epoch are weighted based on the model’s confidence in their prediction (harder examples, with lower confidence, receive higher weights).

We experimented with several other curriculum learning strategies (Supplementary Figure S6): (1) No CL -Curriculum learning was not applied. A user-independent model was fine-tuned on calibration trials. (2) Validation Adaptation CL - used 50% of the test user’s data as validation trials for the model to adapt to the specific user. (3) User-Aware Difficulty Scheduler - used the prior knowledge of difficulty (bad performing users vs good performing users based on user-dependent accuracy) to train the model with curriculum scheduling, i.e., training with good users’ trials first and gradually introducing difficult trials (by adding bad users’ trials). (4) Static Sharded CL: Divided the training data into different shards based on the difficulty metric (variance). Then trained on more similar data more often. (5) Personalized Adaptation CL: Here, the training data was mapped to the calibration data distribution (or test users’ data). Heavier weightage was given to calibration trials for data augmentation, and the difficulty was calculated based on the similarity to calibration trials, and then the curriculum is scheduled, gradually starting from more similar trials to less similar trials.

##### Data Augmentation

To improve robustness and combat overfitting to specific users, we employed two real-time data augmentation strategies during training:

- Gaussian Noise Injection: Randomly added zero-mean noise (*σ* = 0.05) to the MxG input signals with a 50% probability.
- Channel Dropout: With a 30% chance, 10% of channels were randomly zeroed out to simulate sensor noise and force redundancy learning.

##### Training and Evaluation

We employed leave-one-subject-out (LOSO) cross-validation across the users. In each fold, data from one user is completely held out for testing, and the remaining data from other users is used for general training and validation. For calibration-based user adaptation experiments, a few trials (ranging from 1 to 20 per class, randomly selected) were taken from the held-out user and added to the training set. The remaining data from the held-out user was used as the final test set. For each number of calibration trials, a new model was trained with the combined other users’ training data, with added calibration trials.

The model was trained using the AdamW optimizer, with an initial learning rate of 1*e −* 3, weight decay of 1*e* − 4, and a OneCycleLR scheduler for dynamic learning rate adjustment (with max-lr = 3*e*− 3 and pct-start = 0.3). A ReduceLROnPlateau scheduler was also employed, monitoring validation accuracy to further reduce the learning rate if performance stagnated (factor 0.5, patience 10). To accelerate training and reduce GPU memory consumption, we enabled mixed-precision training using PyTorch’s AMP module (torch.amp.autocast). Training was conducted for a maximum of 250 epochs per fold, and the best model checkpoint was selected based on the highest validation accuracy. Performance was evaluated using overall test accuracy on the remaining data from the held-out user.

**Table 3:** Pipeline Details.

| Block | Layers | Configurations | Output Shape |
| --- | --- | --- | --- |
| **Input Block** | Conv1D, Batch-Norm1D, GELU, AvgPool1D, Conv1D (1x1), BatchNorm1D, Residual | Conv1D: 64 channels, kernel=15, padding=7; AvgPool1D: pool size=4; Residual connection added | [B, 64, T/4] |
| **Depthwise Separable Block 1** | DepthwiseConv1D, PointwiseConv1D, SE Block, Max-Pool1D | DepthwiseConv1D: kernel=15; SE: reduction ratio=8; MaxPool1D: pool size=2 | [B, 128, T/8] |
| **Depthwise Separable Block 2** | DepthwiseConv1D, PointwiseConv1D, SE Block, AdaptiveAvgPool1D | DepthwiseConv1D: kernel=9; SE: reduction ratio=8; AdaptiveAvg-Pool1D: output size=25 | [B, 256, 25] |
| **Multi-scale Attention Block** | Linear (QKV), Scaled Dot-Product Attention, Temporal Mean Pool | Projects Q/K/V from [B, 256, 25]; computes attention; pooled to [B, 256] | [B, 256] |
| **Classifier Block** | Linear, Spectral-Norm, LayerNorm, GELU, Dropout, Linear | Linear: 256→512 with Spectral-Norm; Dropout: p=0.5; Final Linear: 512→NumClasses | [B, NumClasses] |
| **Loss Function** | LabelSmoothing CrossEntropy | Smoothing factor = 0.1 | - |
| **Regularization** | Dropout, Spectral-Norm | Dropout: p=0.5; SpectralNorm on Linear layers (first linear layer in classifier) | - |
| **Data Augmentation** | Gaussian Noise Injection, Channel Dropout | Noise std=$\sigma = 0.05$ (50% chance); Dropout: zero 10% channels randomly (30% chance) | - |
| **Curriculum Learning** | Dynamic Difficulty Scheduler, Weighted Sampling | Adjusts sample weights & batches based on epoch progress, validation performance, trend, and class balance | - |

## Acknowledgements

The authors would like to thank the participants of the study for volunteering their time.

## Supplementary Results

**Supplementary Figure S1:**
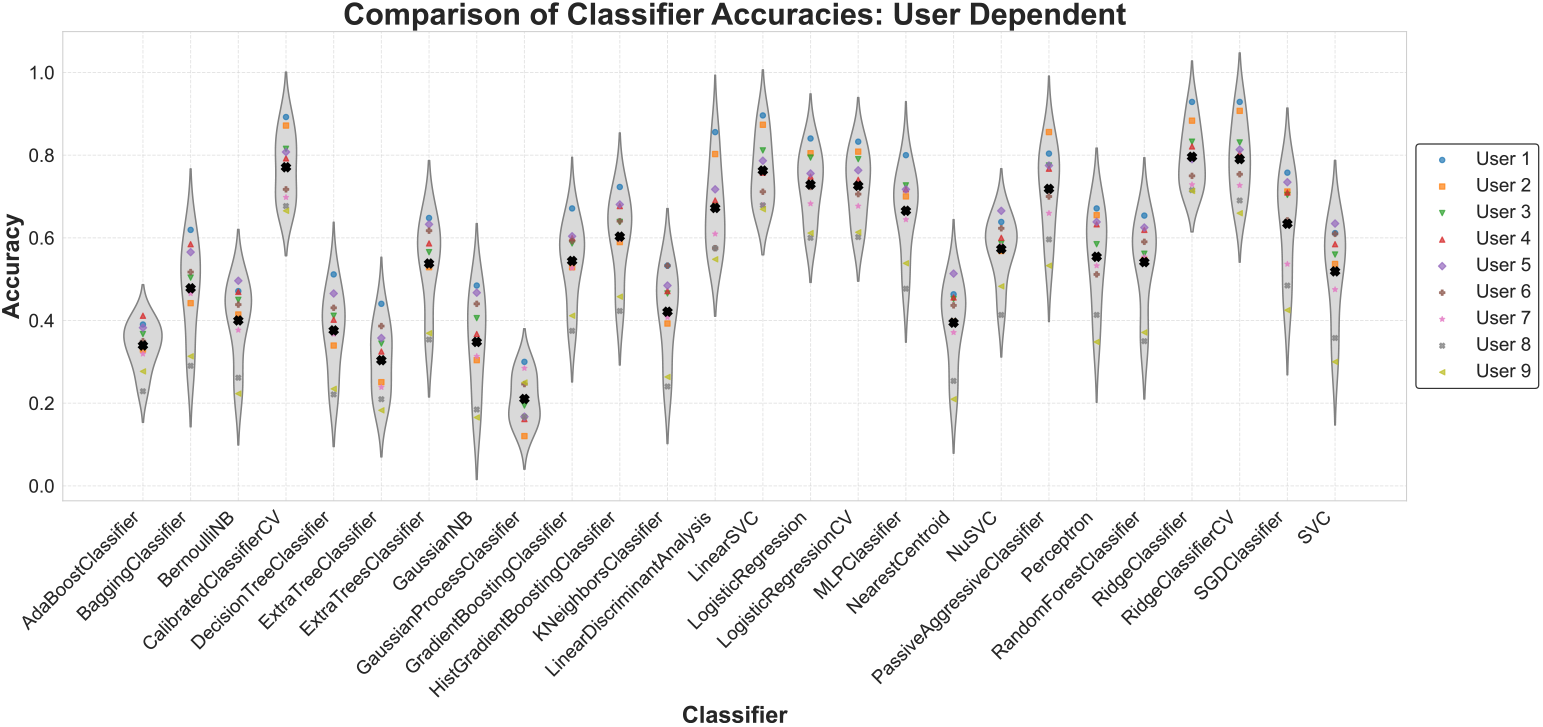
Comparison of classifiers for user-dependent SSR with headphone form factor. The performance distribution of 9 users with 26 different classifiers for user-dependent SSR is shown. Note, Ridge Classifier resulted in the best overall accuracy, which was selected for further analysis.

**Supplementary Figure S2:**
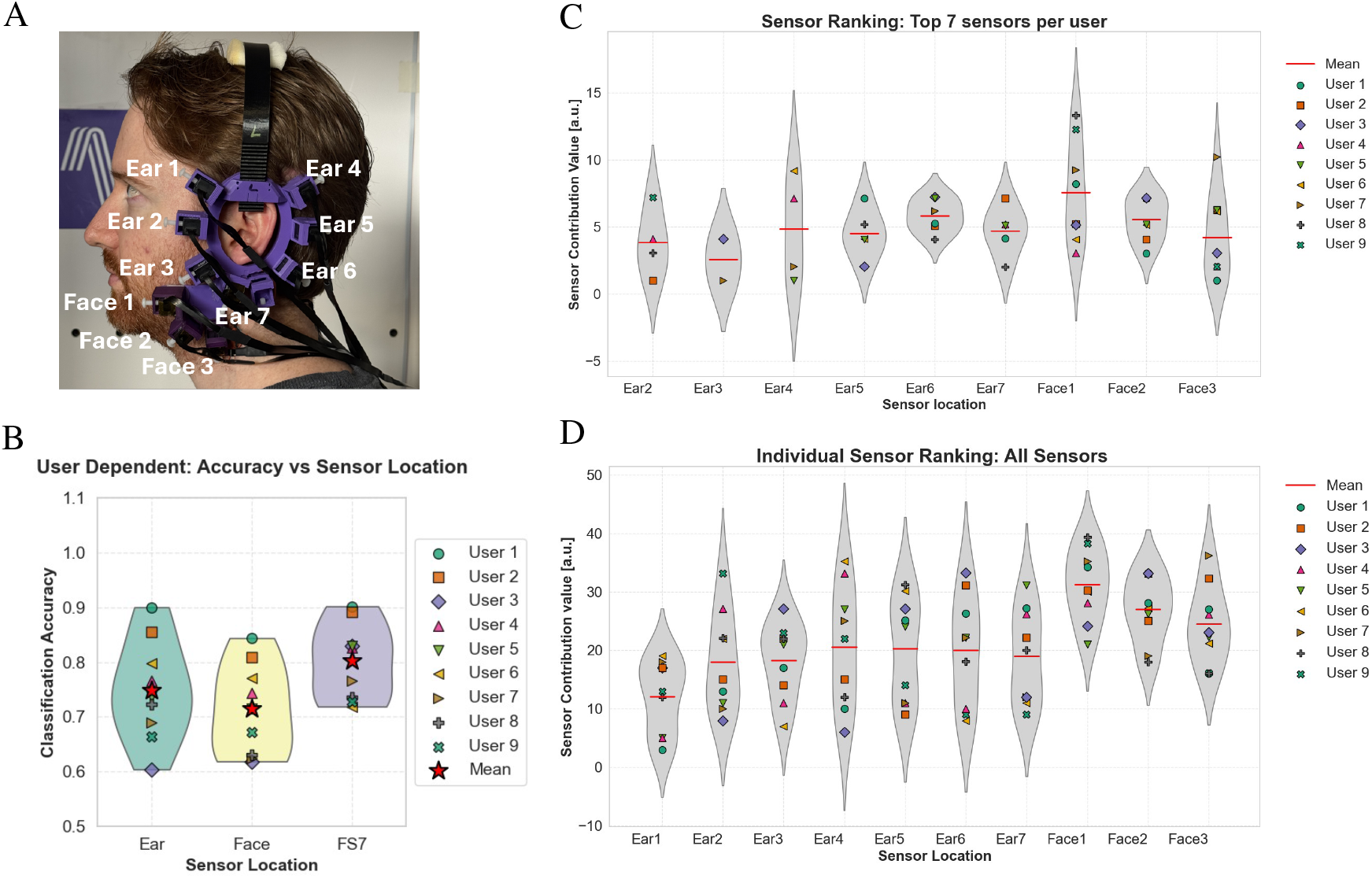
Comparison of SSR performance with different sensor locations. **(A) Sensor location names in the headphone form factor**. Headphone worn by one of the authors for demonstration. 7 sensors were around each side of the ear were named from Ear 1 to Ear 7, 3 of which were at the front, 3 were at the back, and one was below the ear. Three Face sensors were named Face 1, Face 2, and Face 3. **(B) Accuracy comparison of ear and face sensors and of the forward selection strategy**. The performance distribution of 9 users for user-dependent SSR is shown for sensors around the ear, face, and forward selection-based sensor selection with the top 7 sensors. Note, the headphone form factor had 7 sensors around each ear and 3 sensors on each side of the face. FS7 here is the performance with the Forward selection strategy using 7 optimal sensors (from all 20 sensors). The mean accuracy with sensors around the ear was 74.87% and for face it was 71.45%, both significantly lower than FS7 (forward selection strategy with 7 optimal sensors). Note, the lower accuracy for face sensors compared to ear sensors could be due to the lower number of sensors (7 ear sensors vs 3 face sensors at each side) **(C) The location of the** 7 **sensors per user, selected with forward selection**. The distribution of the sensor contribution with the forward selection strategy for each user is shown. Sensor contribution was calculated based on their accuracy contribution and ordering sequence in the forward selection scheme. Note, the Ear 1 sensor was not in the top 7 list. The importance of sensors varied across subjects; however, face sensors and the sensors at the back of the ear, in general, showed higher contribution individually. **(D) Ranking of All Sensors**. The distribution of sensor contribution value for user-dependent SSR for each user is shown. Each marker represents the sensor contribution value (calculated based on their ordering and accuracy contribution in the forward selection scheme) for each user for the particular sensor shown on the x-axis. Note that, when taking all sensors into consideration, individually, face sensors, as expected, had a higher mean contribution to accuracy compared to each individual ear sensors.

**Supplementary Figure S3:**
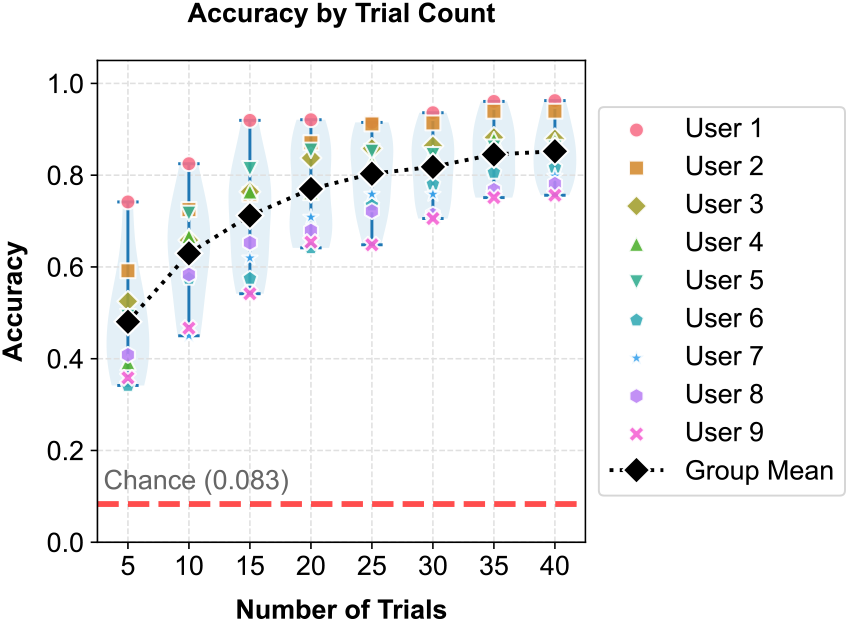
Data Dependency. Performance distribution of 9 users for user-dependent SSR is shown as a function of increasing number of trials (5×). Note, the accuracy increased with more data, as expected, indicating that more trials could have further improved performance.

**Supplementary Figure S4:**
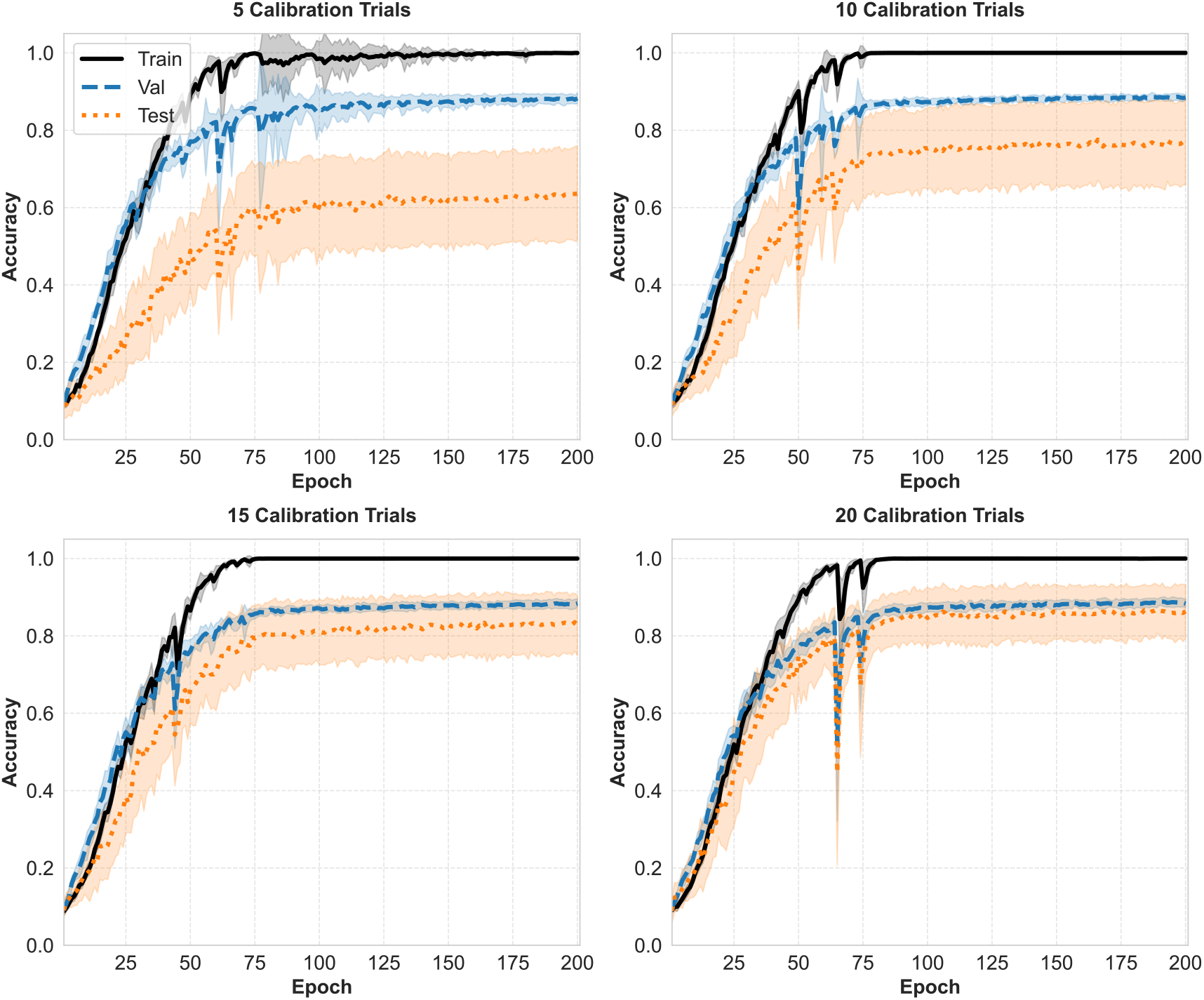
User-independent model learning curves with increasing calibration trials. The learning curves of the user-independent model with averaged training, validation, and test accuracies per epoch for calibration trials 5, 10, 15, and 20 are shown. The corresponding shades show the distribution across users. Notice the reduction in difference between train and validation performance with increasing number of calibration trials, indicating that the model is able to generalize well to unseen data with increased user adaptation

**Supplementary Figure S5:**
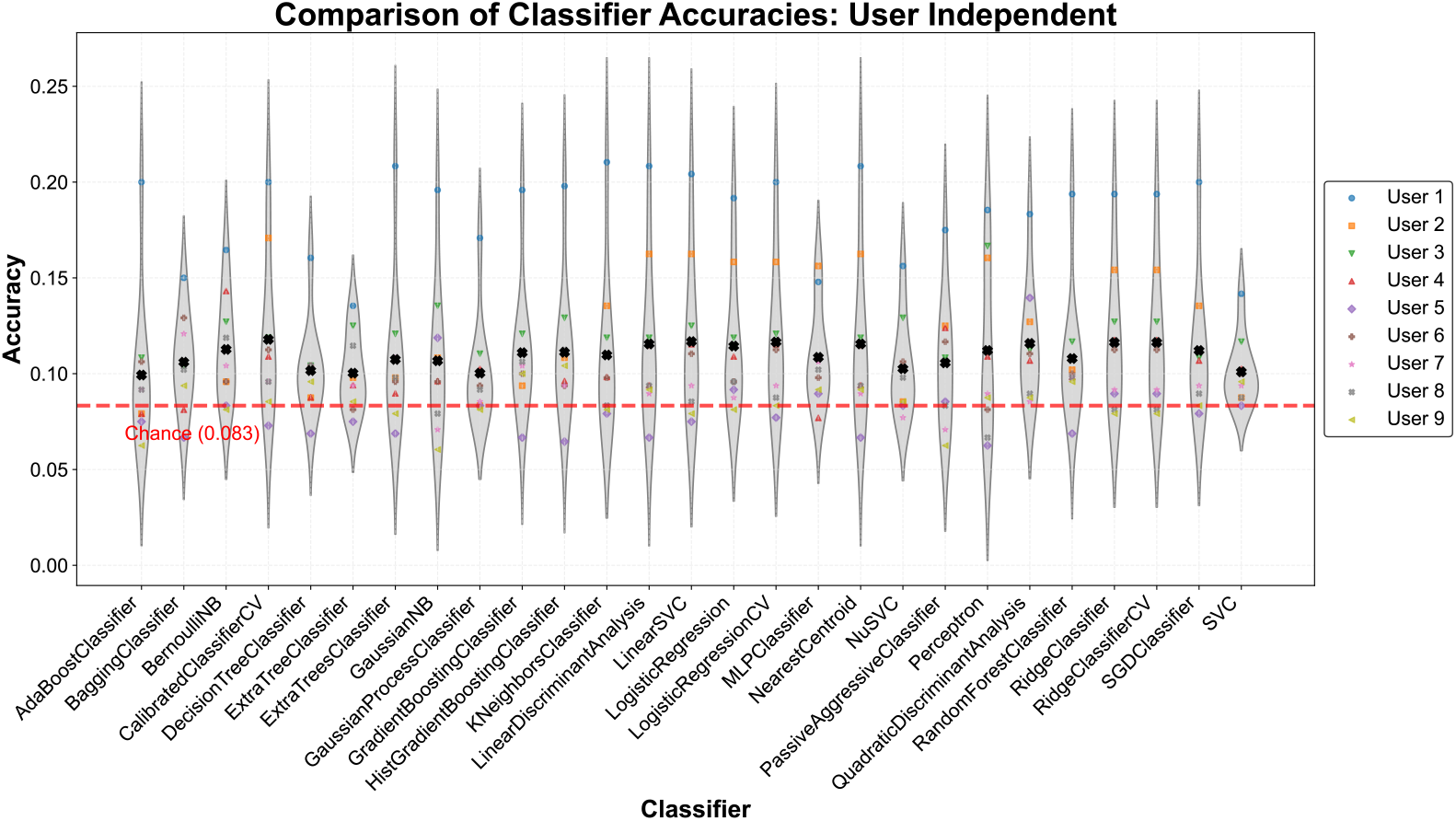
User-independent SSR accuracy with linear models. The performance distribution of the 9 users with 26 different linear classifiers for user-independent SSR is shown. Note that accuracy with most classifiers was just above chance, indicating linear models might be unsuitable to capture generalizability in this task.

**Supplementary Figure S6:**
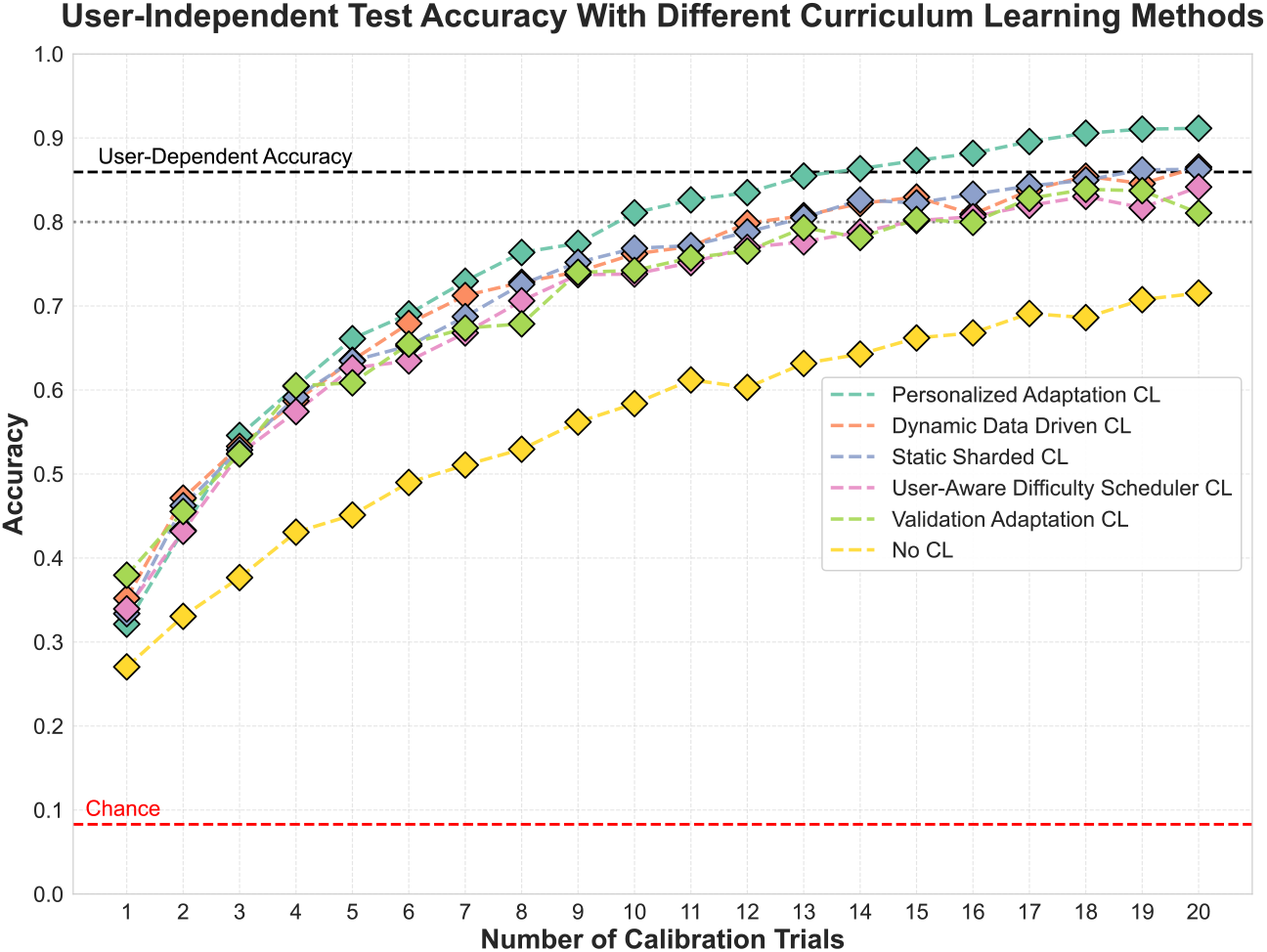
User-independent SSR accuracy with different types of curriculum learning strategy. **No CL**: Curriculum learning not applied. A user-independent model is fine-tuned on calibration trials. **Validation Adaptation CL** - Uses 50% of the test users’ data as validation trials for the model to adapt to the specific test user. **User-Aware Difficulty Scheduler**: Uses the prior knowledge of difficulty (bad subjects vs good subjects based on user-dependent accuracy) to train the model with curriculum scheduling, i.e., train with “good” subjects’ trials first and gradually introduce difficulty (add bad subjects’ trials). **Static Sharded CL**: Divides the training data into different shards based on a difficulty metric (variance). Then trains on more similar data more often. **Dynamic Data-Driven CL**: In this approach, the “difficulty” of the curriculum is calculated dynamically during training based on the model’s performance metrics (like validation accuracy, loss history, and performance trend) for each subject. **Personalized Adaptation CL:** Here, the training data is mapped to the calibration data distribution (or test users’ data). Heavier weightage to calibration trials for data augmentation, difficulty is calculated based on the similarity to calibration trials, and then the curriculum is scheduled gradually, starting from more similar trials to less similar trials.

**Supplementary Figure S7:**
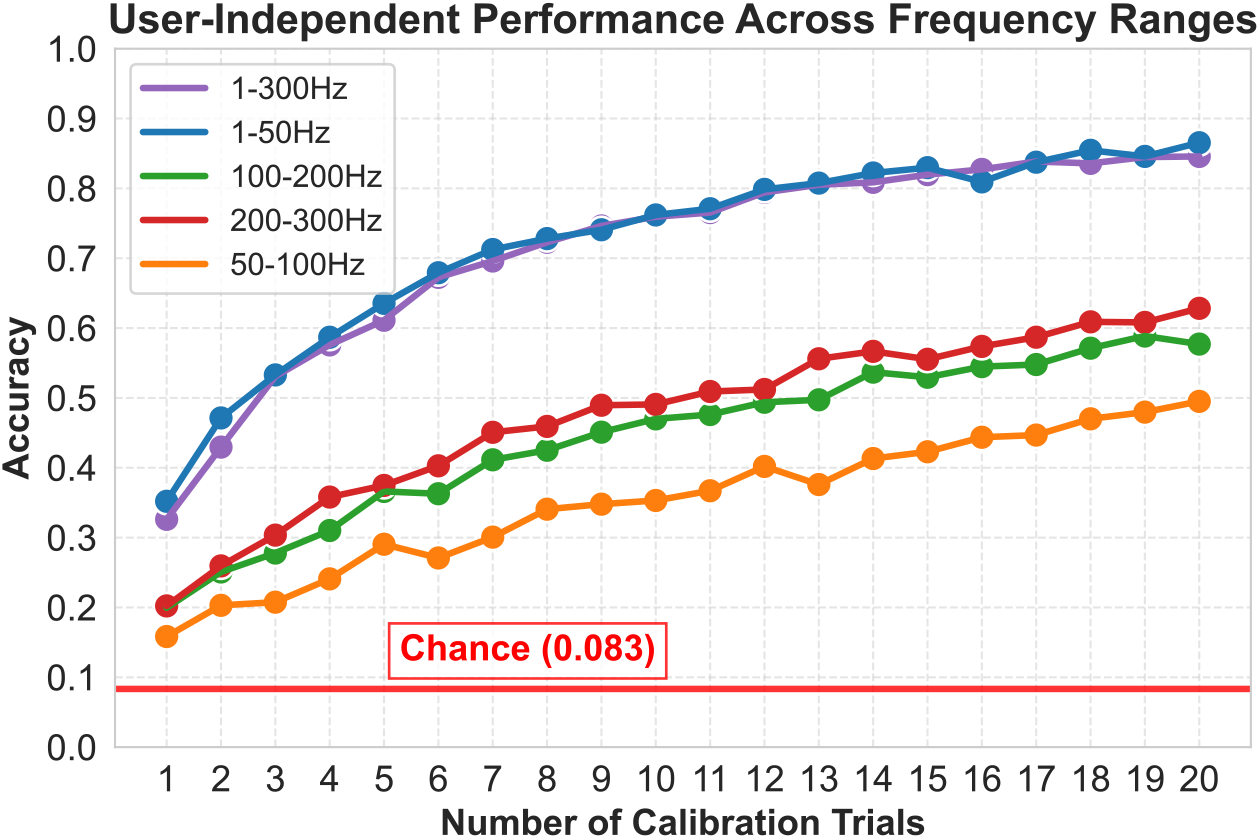
Frequency contributions for user-independent SSR with headphone form factor. The performance trend with increasing number of calibration trials is shown with 5 different frequency ranges: 1 *−* 50 Hz, 50 *−* 100 Hz, 100 *−* 200 Hz, 200 *−* 300 Hz, 1*−* 300 Hz. Similar to user-dependent performance, 1*−*50 Hz and 1 *−* 300 Hz frequency ranges of MxG signals resulted in higher accuracy. All the frequency ranges (including high frequency MxG) showed significantly higher performance (50 − 60%) than chance.

